## supplemental figures and tables for "MicroRNA-deficient embryonic stem cells acquire a functional Interferon response"

**Supplementary Figure 1**

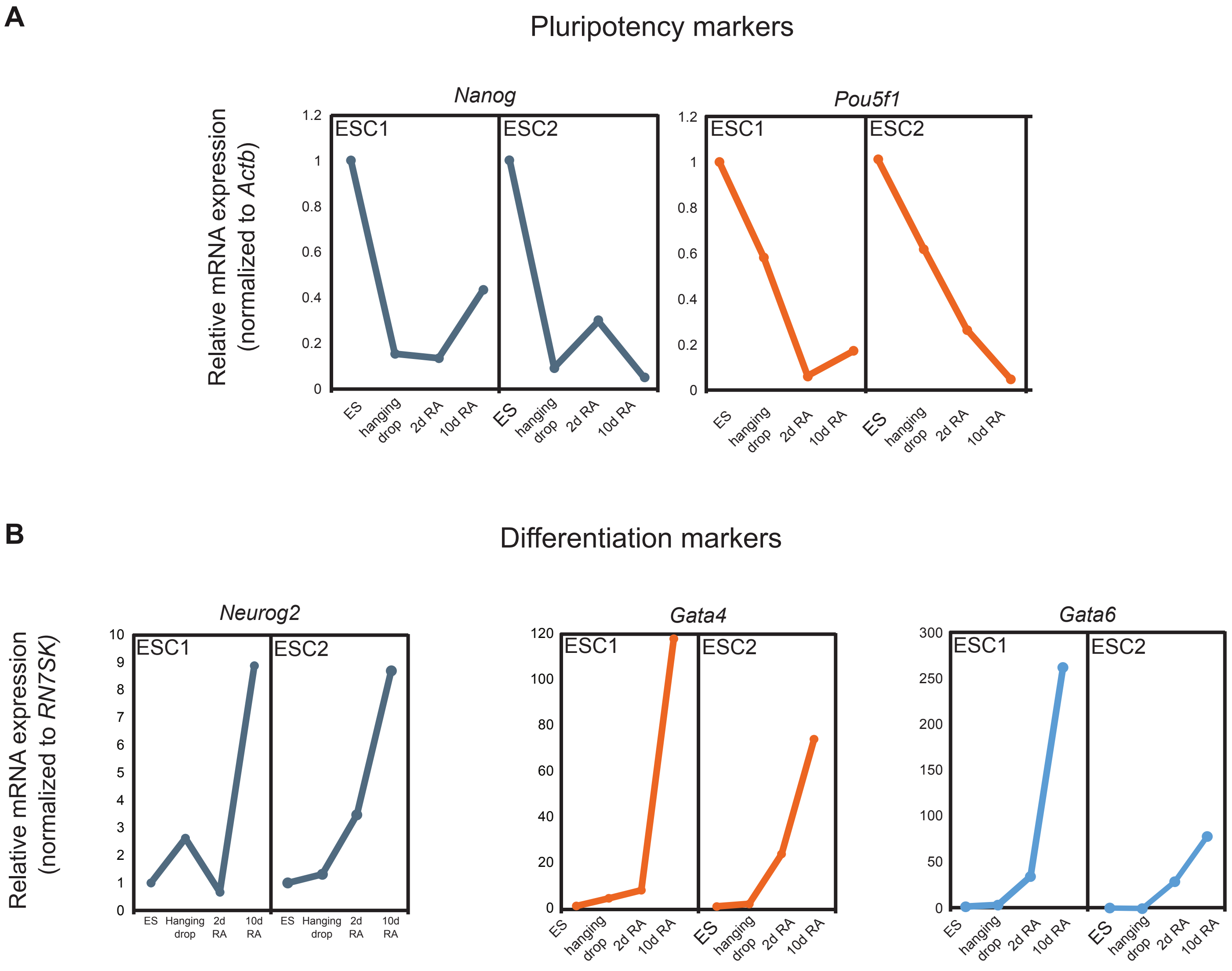


**Supplementary Figure 1, related to Figure 1. Retinoic acid differentiation of ESCs.** (**a**) Quantification of pluripotency markers, *Nanog* and *Pou5f1* (*Oct-4*) expression by qRT-PCR upon differentiation with retinoic acid (RA), and normalized to *Actb* expression, during pluripotency (ESC), after 2 days in hanging drops, 2 days (2d RA) and 10 days in retinoic acid (10d RA). (**b**) Quantification of differentiation markers expression, *Neurog2*, *Gata6* and *Gata4*, during a time course differentation with retinoic acid, as in (**a**). Both panels include data of one representative experiment.

### Supplementary Figure 2

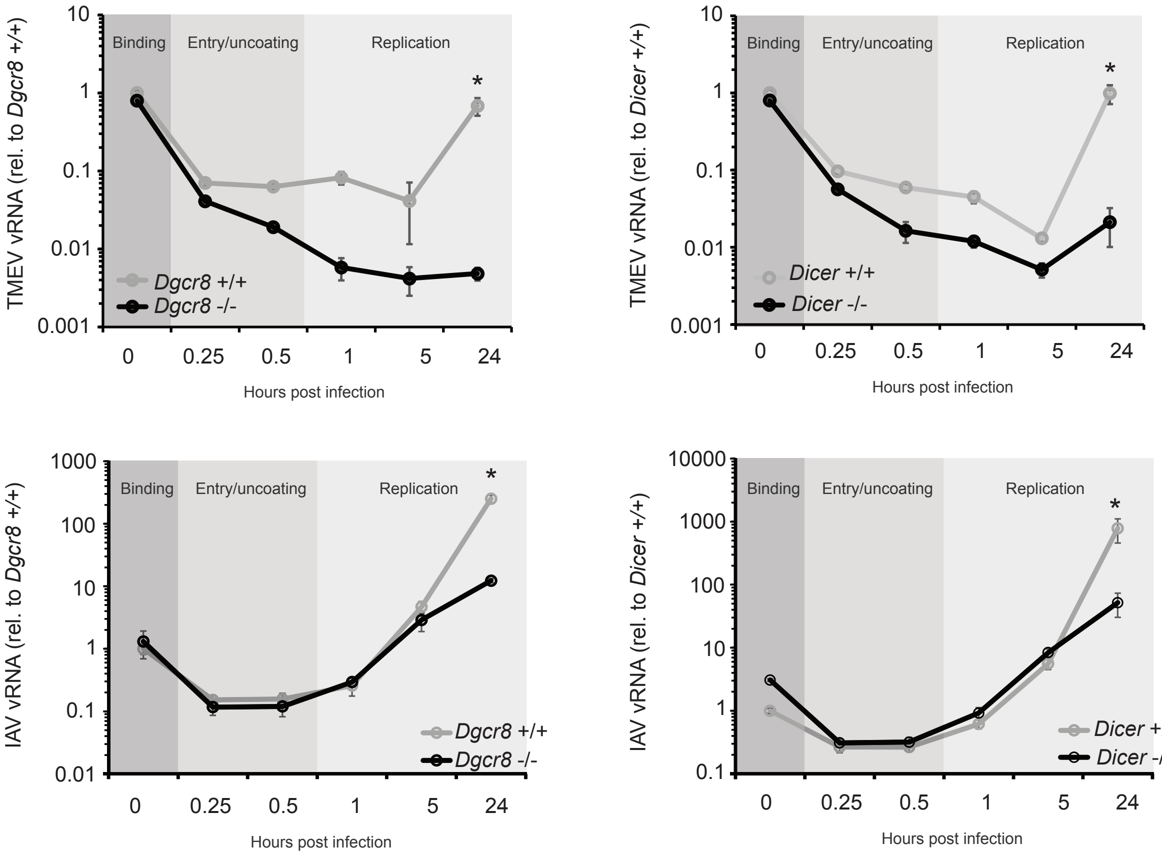


**Supplementary Figure 2, related to Figure 2. Viral entry in miRNA-deficient and wild-type ESCs**
To rule out morphological changes affecting the outcome of viral infections, a binding and entry assay was performed. Cells were infected at 4°C which inhibits entry of viral particles, but allows binding of viruses. At t=0, cells are moved to 37°C resulting in a synchronized entry of virus particles and allowing an accurate quantification of how much virus enters the cell and virus replication kinetics. Cells were infected with either TMEV (top panels) or Influenza A virus (bottom panels) and viral RNA was quantified by qRT-PCR. For all viruses, binding was comparable in all cell lines. Differences in TMEV kinetics were greater soon after the entry phase and continued to increase towards the end of the experiment. Differences in IAV became most apparent late in the replication phase, possibly because IAV is very proficient in knock down the IFN response in its host cells. Data are average (n=3) +/- s.d, (*) denotes p-value<0.05 by t-test.

### Supplementary Figure 3

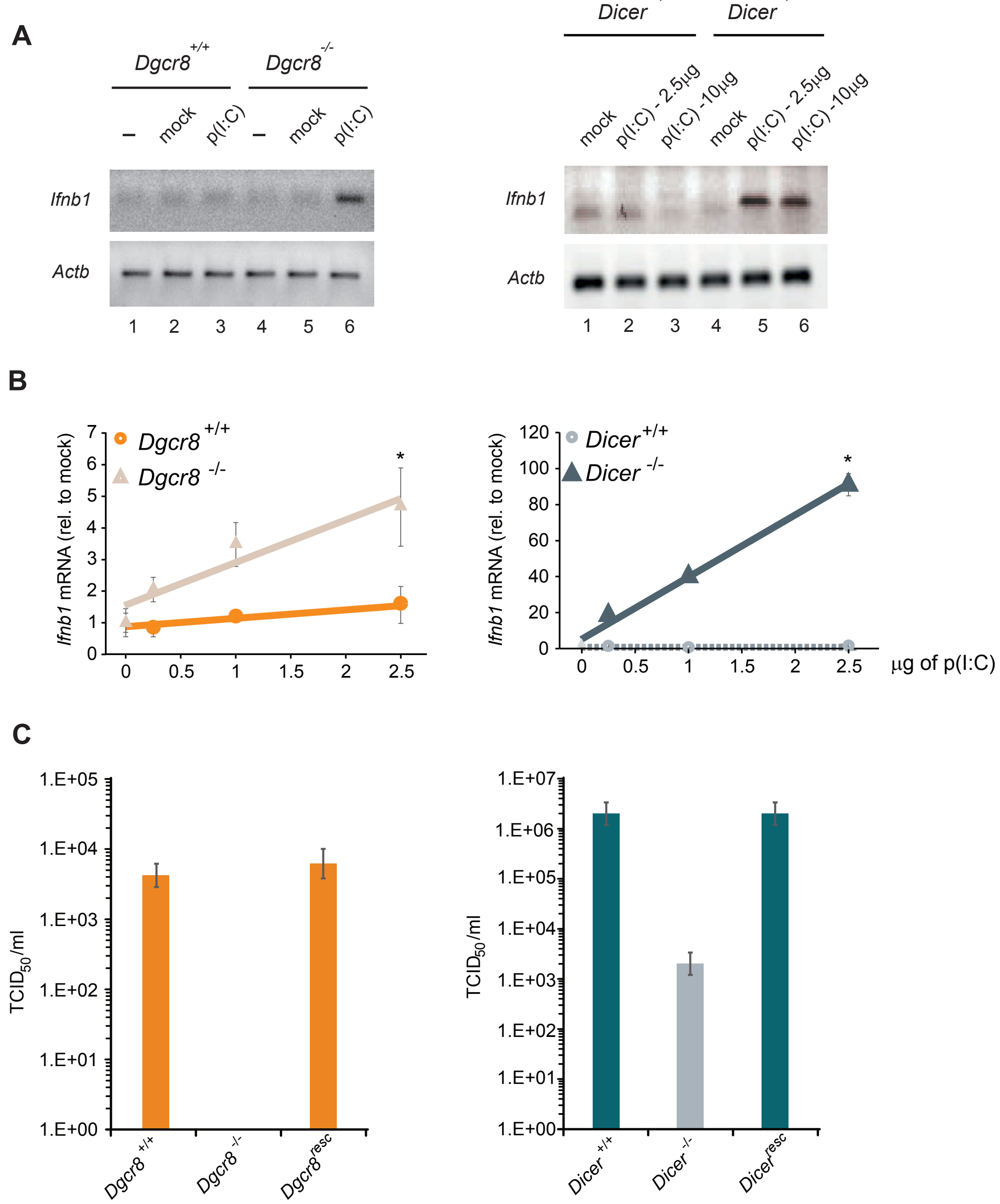


**Supplementary Figure 3, related to Figure 2. miRNA-deficient ESCs acquire IFN expression in response to dsRNA (poly(I:C)).**(**a**) RT-PCR of poly(I:C) activated ESCs. Only *Dgcr8 -/-* (lane 6, left panel) and *Dicer -/-* (lane 5 and 6, right panel) express detectable *Ifnb1* transcript (top panels). *Actb* amplification serves as a loading control. (**b**) *Ifnb* mRNA expression after increasing amounts of poly(I:C) in *Dgcr8-/-* and *Dicer-/-* mESCs. Average (n=3) +/- sd, normalized to mock, (*) p-value <0.05 by t-test. (**c**) TCID50 quantification after TMEV infection of *Dgcr8* (left) and *Dicer* (right) rescued cell lines.

### Supplementary Figure 4

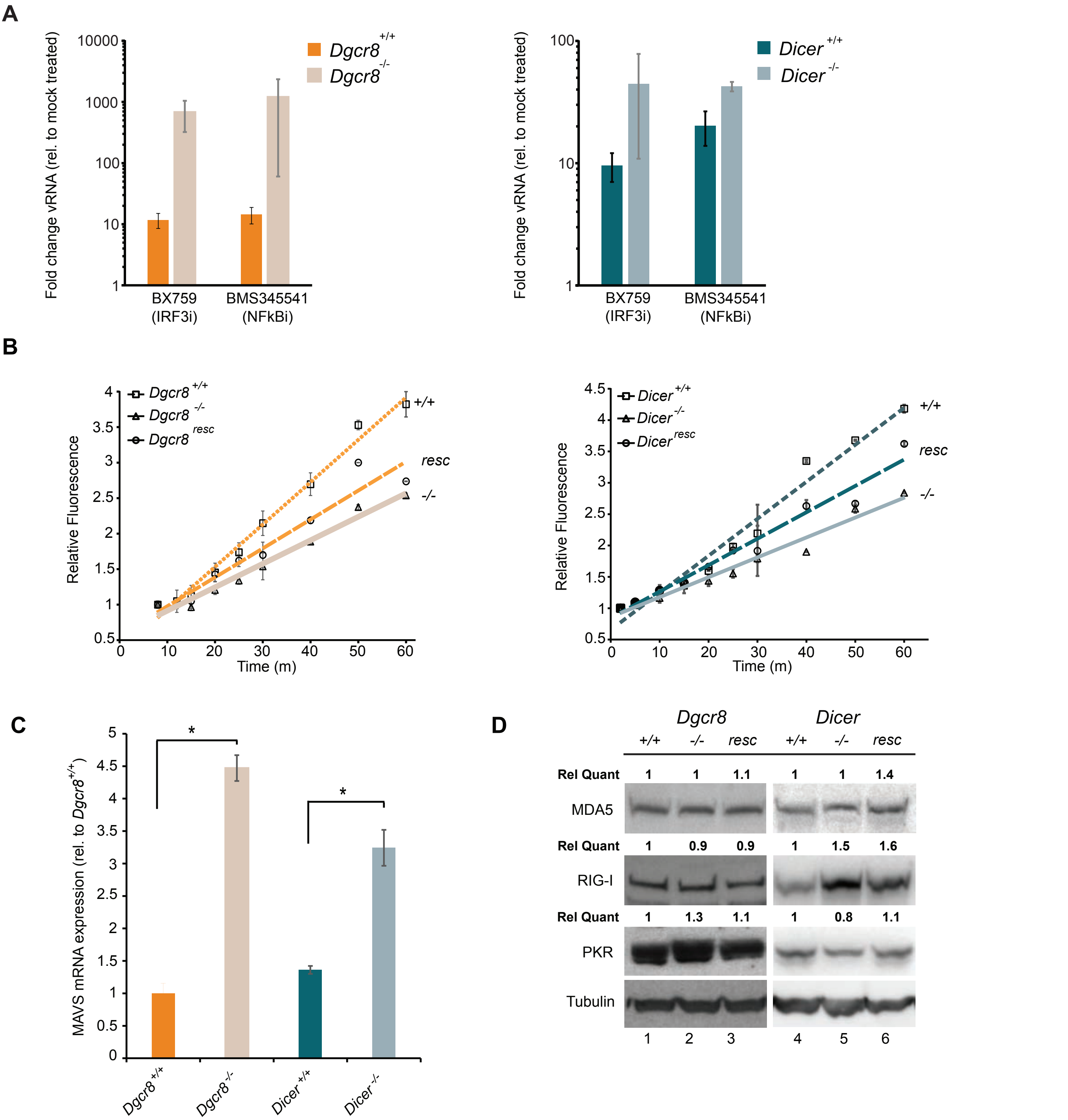


**Supplementary Figure 4, related to Figure 3. miRNAs regulate MAVS and mitochondrial activity**
(**a**) Treatment of mESCs with inhibitors of IRF3 and NF-κB activation improves viral replication to a much greater extend in the *Dgcr8-/-* and *Dicer-/-* cells compared to the parental cells, suggesting that the inhibitory effect that miRNAs have on the IFN response is upstream of these two transcription factors. (**b**) Quantification of oxidative phosphorylation activity by Rhodamine 123 uptake assay in *Dgcr8* (left) and *Dicer* (right) cell lines. (**c**) qRT-PCR quantification of MAVS mRNA. MAVS is higher expressed in the absence of miRNAs (*Dgcr8-/-* and *Dicer-/-*), when compared to parental cell lines (*Dgcr8+/+* and *Dicer+/+)*. Data show average (n=3)+/- s.d, (*) denotes $p<0.05$, t-test. (**d**) Western blot analyses for MDA5, RIG-I and PKR for wild-type cell lines (+/+), *Dgcr8 and Dicer* KO (-/-), and the respective rescued cell lines (resc). Quantification of the signal normalized to Tubulin and relative to wild-type is shown at the top of each panel.

### Supplementary Figure 5

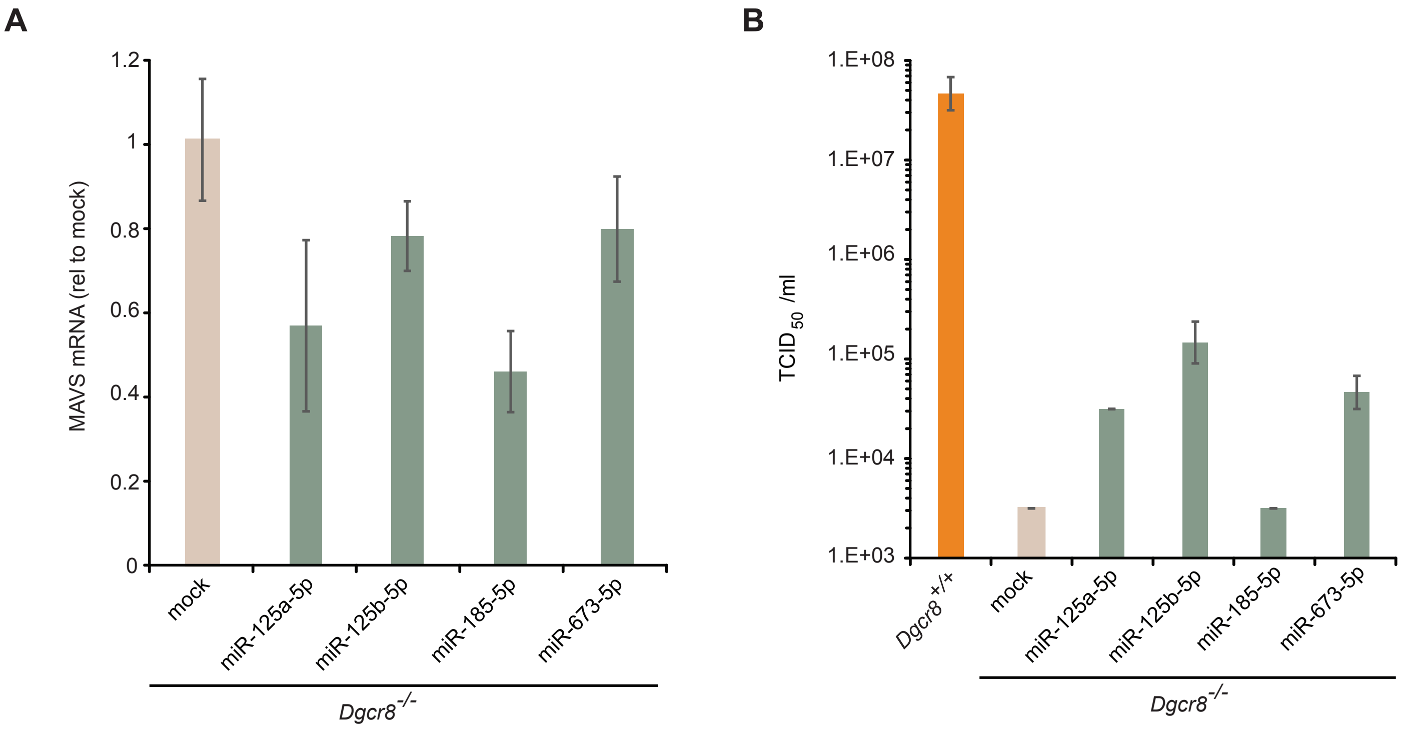


**Supplementary Figure 5, related to Figure 5. Characterization of MAVS expression in ESCs and regulation by miRNAs**

(**a**) *Dgcr8-/-* cells were transfected with miRNA mimics for 24 hours, followed by RNA extraction and MAVS mRNA quantification by qRT-PCR. Transfection of all miRNA mimics result in (non-significant) decrease of MAVS levels. (**b**) *Dgcr8-/-* mESCs were first transfected with mimics, as in (**a**) followed by TMEV infections 36-hours later, and measurement of the TCID50. All mimics except for mir-185 resulted in higher TCID50 values compared to mock transfected cells, suggesting that in mESC cells targeting MAVS with these miRNAs leads to a higher susceptibility to viral infection.

### Supplementary Figure 6

**A**

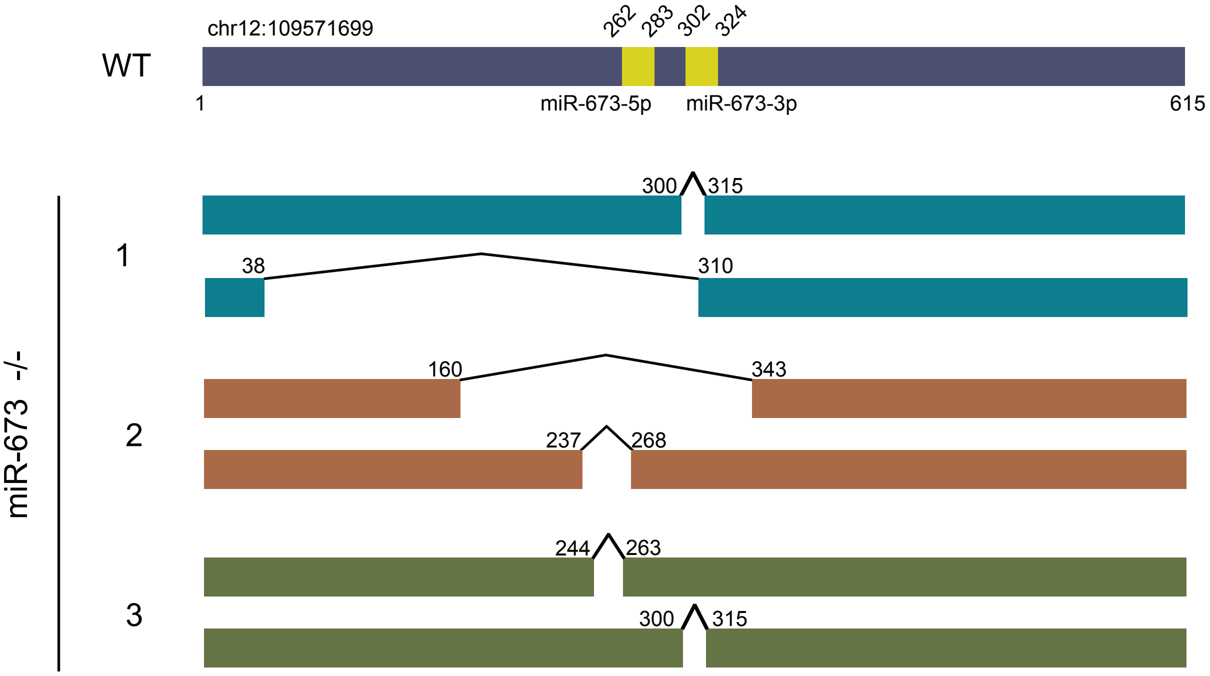


**B**

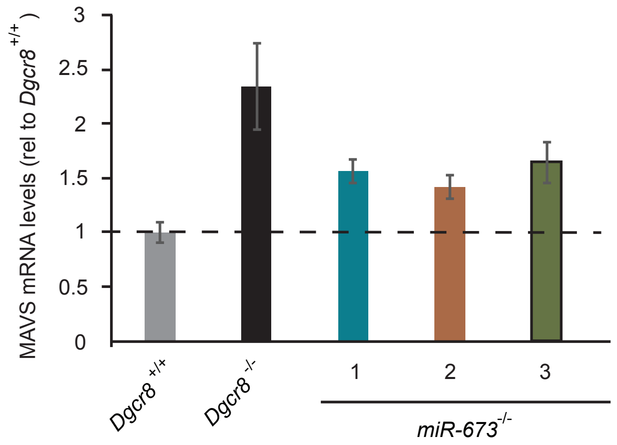


**Supplementary Figure 6, related to Figure 5. Characterization of miR-673 negative mouse embryonic stem cells.**

(**a**) Genomic sequencing of miR-673 locus contained in chromosome 12. In WT, the position of the two miRNAs encoded by the locus, miR-673-5p and miR-673-3p are indicated by yellow boxes. For miR-673-/- clones (1,2 and 3) both sequenced alleles are shown with the indicated deletions induced by CRISPR. (**b**) Quantification of mmu-miR-673-5p abundance by qRT-PCR. Data shows the average (n=3) +/- s.d, normalized to U6 snRNA. (n.d, not detected) (**c**) Quantification of MAVS mRNA levels by RT-qPCR. Data show the average of at least (n=3)+/- s.d, normalized to *Actb,* and relative to *Dgcr8 +/+.*

### Table S1. Oligonucleotides used in this study

| | Name | Sequence (5'-3') |
|---|---|---|
| **qRT-PCR** | MAVS forward | CTGCCTCACAGCTAGTGACC |
| | MAVS reverse | CCGGCGCTGGAGATTATTG |
| | RIG-I forward | CACTTCGTTCATCTCTGGCG |
| | RIG-I reverse | AGAGTGAGGCAGCTTCCATT |
| | MDA5 forward | TCATCGAAGCAGCTGACACT |
| | MDA5 reverse | GCCTGGAACGTAGACGACAT |
| | TMEV forward | TGTGGACTTGGACGATGACG |
| | TMEV reverse | CAGTATCGCATACGAGCGGT |
| | Influenza A virus (seg. 5) forward | ATCATGGCGTCTCAAGGCAC |
| | Influenza A virus (seg. 5) reverse | CCGACGGATGCTCTGATTTC |
| | IFNB1 forward | AAGAGTTACACTGCCTTTGCCATC |
| | IFNB1 reverse | CACTGTCTGCTGGTGGAGTTCATC |
| | Dgcr8 forward | GCTGCAGGAGTAAGGACAGG |
| | Dgcr8 reverse | TCGAGCACTGCATACTCCAC |
| | Dgrc8 (exon 3) forward | TCCCAAGAAGAGGCGAATGG |
| | Dgrc8 (exon 3) reverse | CCACGACTTTTGAGCACTGTTT |
| | Dicer forward | GGTCCTTTCTTTGGACTGCCA |
| | Dicer reverse | GCGATGAACGTCTTCCCTGA |
| | GAPDH forward | TGTGTCCGTCGTGGATCTGA |
| | GAPDH reverse | CCTGCTTCACCACCTTCTTGA |
| | Actb forward | GGCACCACACCTTCTACAATG |
| | Actb reverse | GGGGTGTTGAAGGTCTCAAAC |
| | 7SK forward | GACATCTGTCACCCCATTGA |
| | 7SK reverse | GCCTCATTTGGATGTGTCTG |
| | Nanog forward | AGGGTCTGCTACTGAGATGCTCTG |
| | Nanog reverse | CAACCACTGGTTTTTCTGCCACCG |
| | Pou5f1 forward | AGTTGGCGTGGAGACTTTGC |
| | Pou5f1 reverse | CAGGGCTTTCATGTCCTGG |
| | Neurog2 forward | GACATTCCCGGACACACAC |
| | Neurog2 reverse | CCAGCAGCATCAGTACCTCC |
| | Gata4 forward | GAAAACGGAAGCCCAAGAACC |
| | Gata4 reverse | TGCTGTGCCCATAGTGAGATGAC |
| | Gata 6 forward | GCAATGCATGCGGTCTCTAC |
| | Gata 6 reverse | CTCTTGGTAGCACCAGCTCA |
| **RT-PCR** | IFNB1 forward | AGCTCCAAGAAAGGACGAACAT |
| | IFNB1 reverse | GCCCTGTAGGTGAGGTTGATCT |
| **Cloning** | MAVS pLenti FW EcoRV | GCCGGGATATCATGACATTTGCTGAGGACAAGAC |
| | MAVS pLenti RV XbaI | CGGCCCTCTAGATCACTGGGCCAGGCGCCTAC |
| | DGCR8 pLenti-FW EcoRV | TCTAGTCCGATATCATGGAGACAGATGAGAGCCC |
| | DGCR8 pLenti-RV Xba1 | CTGATCACTCTAGATCACACGTCCACGGTGCACAG |
| **3'UTR** | RIG-I_3'UTR_FW_XhoI | GCGCGCGCTCGAGCCTCAGGCTTCTCCGTCTCGTG |
| | RIG-I_3'UTR_RV_NotI | CGGCCCGCGGCCGCAAATACATAAATTTTTAATTAATT |
| | MDA5_3'UTR_FW_XhoI | GCGCGCGCTCGAGCACTTGATTCATGATTATTTTA |
| | MDA5_3'UTR_RV_NotI | CGGCCCGCGGCCGCAATAGTATCTGAAATATACATC |
| | MAVS_3UTR_FW_XhoI | GCGCGCGCTCGAGAGCCTCAGCTGTATGCTGTTCTC |
| | MAVS_3UTR_RV_NotI | CGGCCCGCGGCCGCTTTGCACAGATAAACTCTTTTAATTATCCTG |
| **Probes** | mmu-miR-130a-3p | CAGTGCAATGTTAAAAGGGCAT CCTGTCTC |
| | mmu-miR-293-3p | AGTGCCGCAGAGTTTGTAGTGT CCTGTCTC |
| | mmu-miR-294-3p | AAAGTGCTTCCCTTTTGTGTGT CCTGTCTC |
| **CrRNA target** | 1 | CCAGAGAAAATGTTGCTCCGGGG |
| **mmu-miR-673** | 2 | ACCAGAGCTGTGAGCCCCTCAGG |
| | Genomic PCR miR-673 locus F | AGCAGTGATGGGTGTGCTAC |
| | Genomic PCR miR-673 locus R | TCCATTTCCCATCCCCTTGC |
